# Knowledge of Chagas disease in Latin American migrant population living in Japan and factors associated with knowledge level

**DOI:** 10.1101/589051

**Authors:** Inés María Iglesias-Rodríguez, Shusaku Mizukami, Dao Huy Manh, Tieu Minh Thuan, Hugo Alberto Justiniano, Sachio Miura, George Ito, Nguyen Tien Huy, Kenji Hirayama

## Abstract

**Background:** Chagas disease (CD), typically confined to the Latin America (LA) region, is emerging as a global health problem. In Japan, as in the rest of world, the under-diagnose rate of CD is alarmingly high. Various studies have highlighted the importance of informed knowledge in the seeking behavior. Educational integrative activities, with consideration for socio-cultural factors, can help increase the knowledge of the participants. There has been no studies that analyze the difference in knowledge, before and after these educational activities. This study aimed to qualitatively and quantitatively investigate the knowledge, behavior and attitude toward CD among LA migrants in Japan and to evaluate the effectiveness of the community educational activity in increasing knowledge of CD.

**Methodology:** This cross-sectional study involved two questionnaires to analyze the knowledge of the LA migrant participants before and after the community activity (CA) in four cities in Japan (Oizumi, Suzuka, Hadano, and Nagoya).

**Principal Findings:** A total of 75 participants were enrolled, predominantly Bolivians from hyperendemic areas. The baseline knowledge of CD was low. However, most of them were familiar with the disease although less than 10% of them had been tested for CD before. Living in Japan for more than 10 years and previously being tested for CD were the factors associated with better knowledge. The conducted CA significantly improved the knowledge of the participants. They associated the term “Chagas” mostly with fear and concern. In contrast to other studies, the level of stigmatization was low. The barriers in care seeking behavior were language, migration process and difficulties to access to the healthcare system.

**Conclusion:** Educational activities with integrative approach are useful to increase knowledge of CD. The activity brings the possibility to explore not only the level of knowledge, but also to reveal the experience and to understand the needs of the people at risk.

**Author Summary:** Though the incident rate of Chagas disease (CD) has fallen, more than 7 million people are affected worldwide. The CD prevalence is under-estimated because just 1% of these affected people can access to the diagnosis and treatment. This situation is maintaining mainly for the lack of implication of socio-cultural factors in the interventions to decrease the burden of the disease. Educational activities with integral approach are useful to increase the knowledge of the people at risk. People that have being tested for CD before or living in Japan for more than 10 years have better knowledge about the disease, suggesting the importance of knowledge in the seeking behavior. The authors recommend the implementation of educational activities with integral approach as a strategy to improves the knowledge of Chagas disease among Latin America migrants in Japan.

## Introduction

It has been more than 100 years since the discovery of Chagas disease (CD) and although the number of incident rate has fallen, more than 7 million people are affected worldwide. The prevalence is under-estimated because just 1% of these affected people can access to the diagnosis and treatment [1]. Each year 39,000 new cases occur of which 9,000 results from congenital transmission [2]. In Latin America (LA), CD presents as one of the tropical disease with the highest burden, which is measured in 806,170 disability-adjusted life-years (DALYs). The economic impact is about 627 million dollars in health expenditure worldwide [3].

As a result of increasing migration in the last few years, the disease silently spreads into the United States, Europe and the Western Pacific region [4]. Japan ranks third in the world for migrant population from LA. Brazilian, Peruvian and Bolivian population represent the majority of the Latin American migrants living in Japan. In Japan, there is an estimate of 3000 cases of CD. Nowadays, blood transfusion control is implemented as the only one measure to control CD in Japan [5]. There is an important gap in epidemiological data and cases reports, with most of the population affected undiagnosed [6]

The fact that CD is an emerging disease in non-endemic countries, along with the lack of awareness of the health personnel and its silent course, makes it difficult for detection. In addition to this challenging situation, migrant population are exposed to many barriers to access to the health system.

The migrant population has a disadvantageous situation compared with the native citizens of the of the host country. Language, lack of adaptability of the health system and priorization of their migratory goals in the process of care seeking behavior are described as main barriers. Currently less than 10% of the people affected by CD are diagnosed in non-endemic countries [1,7,8]. Some studies pointed it out that the lack of knowledge of the people at risk have strong influence in the care seeking behavior.

It is important for the population at risk of contracting CD to have the necessary knowledge to be able to combat it through their daily activities. However, to have better outcomes, this knowledge should not be presented as information provided by the scientific community. Such knowledge should be incorporated with the population beliefs and conceptions [9]. Conceptions are the knowledge that each one must explain and situate themselves in their environment. These are result of their history, their environment, their cultural context, their reality and the interactions between all these elements. Thus, the conceptions of who learns should become the starting point of any educational project [10]. Furthermore, we know that the emergence and persistence of CD associated with socio-cultural factors [11]. These are the reasons why educational activities to improve the knowledge should involve an integrative approach and consider the affected people as main actors. However, most of the interventions do not combine all clinical, environmental, sociocultural aspects together. Incorporate the experience of people affected can be an important tool for developing program with better outcomes. An activity presented by the group Conicet “Hablamos de Chagas” is an example of integrative approach where Chagas is considered in all aspects (epidemiological, biomedical, political and sociocultural) [12]. To our knowledge there is no study that analyze the improvement in knowledge of CD after these educational activities.

The primary objectives of our study are to reveal the current knowledge and attitudes towards CD in LA migrant population living in Japan; to analyse the factors associated with better knowledge and to evaluate the effectiveness of a community activity in increasing the knowledge of CD. The secondary objective is to identify possible strategies to overcome existing barriers for the migrant population to access care.

## Methods

### Study design

This cross-sectional study involved both quantitative and qualitative methods. The quantitative part was based in two questionnaires administered pre (first questionnaire) and post (second questionnaire) an educational activity where CD was covered in an integrative approach.

More than half of the questionnaire were based on two previous studies, one conducted in United States to assess the awareness among LA migrant population living in Los Angeles [13] and other in Geneva, Switzerland [14]. The qualitative part was based on one of the educational activities proposed in the book “Hablamos de Chagas” by the Conicet group “Taller I: Construcción colectiva del caleidoscopio”. We decided to modify the videos proposed originally by Conicet group of this activity. Three of the videos of Juana and Mateo: “Salud y Enfermedad”, “Pregnancy” and “Ciudad” were chosen to adapt to the necessities of the migrant population living in non-endemic country [12].

### Study areas and study population

The activity was held from March to June of 2018 in four cities in central Japan:

Oizumi (Gunma prefecture), Hadano (Kanagawa prefecture), Nagoya (Aichi prefecture) and Suzuka (Mie prefecture). They were selected for the high number of LA migrant population. We sought 100 participants. Inclusion criteria were LA migrant adult population (18 years old or more) who are residing in Japan.

### Recruitment method

We collaborated with the Bolivian embassy who worked with the leader of the community to recruit the participants. The activity was also informed by different media directed to Latin-American migrant population living in Japan: radio, magazine and social network. During the day of the activity, the people that came to the embassy for other issues were invited to participate.

### Data collection

After having signed the written informed consent, the first questionnaire was administered to the participant after enrollment. Then the educational activity was conducted. Finally, the second questionnaire including similar questions to the first questionnaire was given. All the communication and education activities with the participants were conducted in Spanish by a native speaker.

The discussion of the educative activity starts with the question “What is the first thing that came to your mind when you hear the word “Chagas”?”. The participants wrote what came in their mind in a piece of paper. It would be pasted in a mural divided in the four dimensions of CD: biomedical, epidemiological, political and socio-cultural. Constructing a kaleidoscopic image of the multidimensionality of CD helps discuss about the problematic in a focal group discussion (FGD) format. Then, some videos of Juana and Mateo was projected and recap of ideas from the videos was conducted [15]. The activity followed with a discussion about barriers that migrant population found in Japan and their suggestion to overcome these problems.

### Variables

The first questionnaire contained 33 questions, 10 of them related to knowledge in the disease. The first questionnaire included questions with regards to demographic information: age, sex, education level, country and city of origin and place of residence. There were questions related to the risk factors of CD: age, living in rural area, living in adobe house, hearing about CD, having a family member who has CD, knowledge of the vector, the risk of being infected by blood transfusion and any information for blood transfusion in LA countries. In addition, there were question related to the risk of transmission: risk of congenital and blood transfusion transmission and risk during the time living in Japan. There were question related to the level of vigilance and policies for CD, in their previous resident areas: living in another country, previously tested for CD and previously treated. There were questions related to knowledge about CD: the magnitude of the problem, severity, routes of transmission, existence of treatment, epidemiology of the disease, symptomatology, difference between disease and positive result of CD diagnostic test and efficacy of the treatment. Questions regarding barriers to access the Health system in Japan and stigmatization were also included.

### Data analysis

The total punctuation was 14.5. To analyse the quantitative data, cross tabulation was conducted for questions not related to knowledge with Stata 2015. For the ones related to knowledge, we calculated the total point of the first questionnaire and divided in 2 groups with a cut off in 60%. To analyse the knowledge of the participants we use R core Team (2018) R. Bivariate analysis was conducted for factors between two groups. The factors with p value less than 0.2 were include in the multivariable analysis. Paired t-test was conducted to analyse if there was improvement after the CA by comparing the total point of people before and after. The FGD was transcribed verbatim in Spanish and translated into English after for publication. Transcription were grouped into categories and by groups to analyse. ArcGIS online was used to create the map that illustrate the Cities of residence of the participants.

### Ethic statements

This study received the approval of the ethics committee of Institute of Tropical Medicine (NEKKEN) of Nagasaki University with the approval number 18031188. Informed written consent was obtained from all participants. All the data collected during the quantitative and qualitative part was anonymized by code number and privacy was protected.

## Results

### Description of the participants

A total of 75 participants were recruited into the study. The participants age ranged between 20 and 70 years old, with a mean of 44.5 ± 13.2 years old. There were slightly more women than men, 54,1% and 45.8%, respectively. Almost 95% of the women had children. Most of the participants had secondary education (62.1%). The Bolivian migrant population represents 95.5% of the participants. More than half of the responders (57,6%) came from endemic areas for CD. The most common city of origin was Santa Cruz in Bolivia (44%), followed by Riberalta (22%). Most of the participants have been living in Japan for more than 10 years (81.1%). Mie prefecture and Kanagawa prefecture had the highest number of participants (Fig. 1). In addition, 9.6% of the participants came from other prefectures including Tokyo (n=8,8 %) and Ishikawa (1.6%) (Table 1).

**Fig 1.**
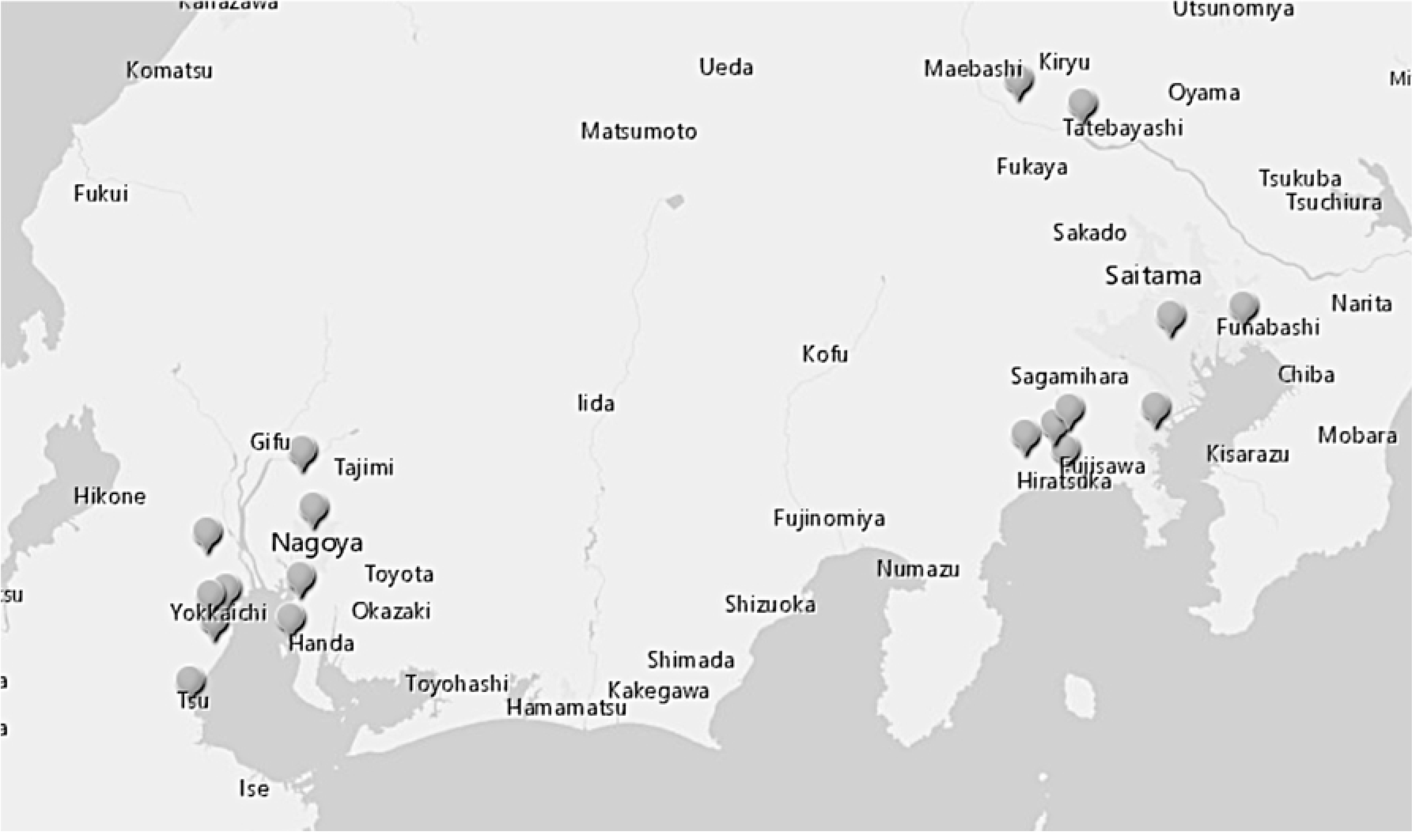
Cities of residence in Japan. ArcGIS online was used to illustrate the Cities of residence of the participants.

**Table 1.** Socio-demographic characteristic of the population

|  | Mean (standard deviation)/number (%) |
| --- | --- |
| **Age (n=64)** | 44.5 (13.2) |
| **Sex** |  |
| - Male (n=33) | 33 (45.8) |
| - Female (n=39) | 39 (54.1) |
| **Education** |  |
| - Primary | 3 (4.5) |
| - Secondary | 41 (62.1) |
| - Universitary | 22 (33.3) |
| **Country of origin** |  |
| - Bolivia | 65 (95.5) |
| - Argentina | 1 (1.4) |
| - Brazil | 1 (1.4) |
| - Japan | 1 (1.4) |
| **City of origin** |  |
| - Hyperendemic | 30 (50.8) |
| - Low endemic | 4 (6.7) |
| - No endemic | 25 (42.3) |
| **Duration living in Japan** |  |
| - 5 years | 9 (13.0) |
| - 5-10 years | 4 (5.8) |
| - 10 years | 56 (81.1) |
| **Resident prefectures in Japan** |  |
| - Gunma | 8 (12.9) |
| - Mie | 25 (40.3) |
| - Kanagawa | 14 (22.5) |
| - Aichi | 9 (14.5) |
| - Tokyo | 5 (8.0) |
| - Ishikawa | 1 (1.6) |
| **Having lived in other countries different to the country of origin.** |  |
| **Non-endemic** |  |
| - Spain | 4 (36.0) |
| - France | 1 (9.1) |
| - USA | 1 (9.1) |
| **Latin America countries** |  |
| - Bolivia | 2 (18.1) |
| - Brazil | 1 (9.1) |
| - Argentina | 1 (9.1) |
| - Brazil & Spain | 1 (9.1) |
| **Female with children** | **36 (94.74)** |
| **People who have donated blood** | **20 (28.2)** |

### Characteristic of the participants related with risk factors of Chagas disease

Most of the responders had heard about CD (82.2%). However, just five responders (7.2%) had been tested for Chagas before. More than half of the responders came from rural areas (60.3%). Nearly a third of the responders have lived in an adobe house (31.67%). Most of the responders did not recognize the vector (63,5%). Out of the people that saw the insect (n=27), 12 of them saw it at home.

In our study, 22.7% of the responders had at least one family members infected by CD. Just two participants referred that their mothers were infected by CD. Regarding blood transmission, 10% of the responders received blood transfusion in LA countries. Nearly 30% of the participants had donated blood previously (Fig. 2).

**Fig 2.**
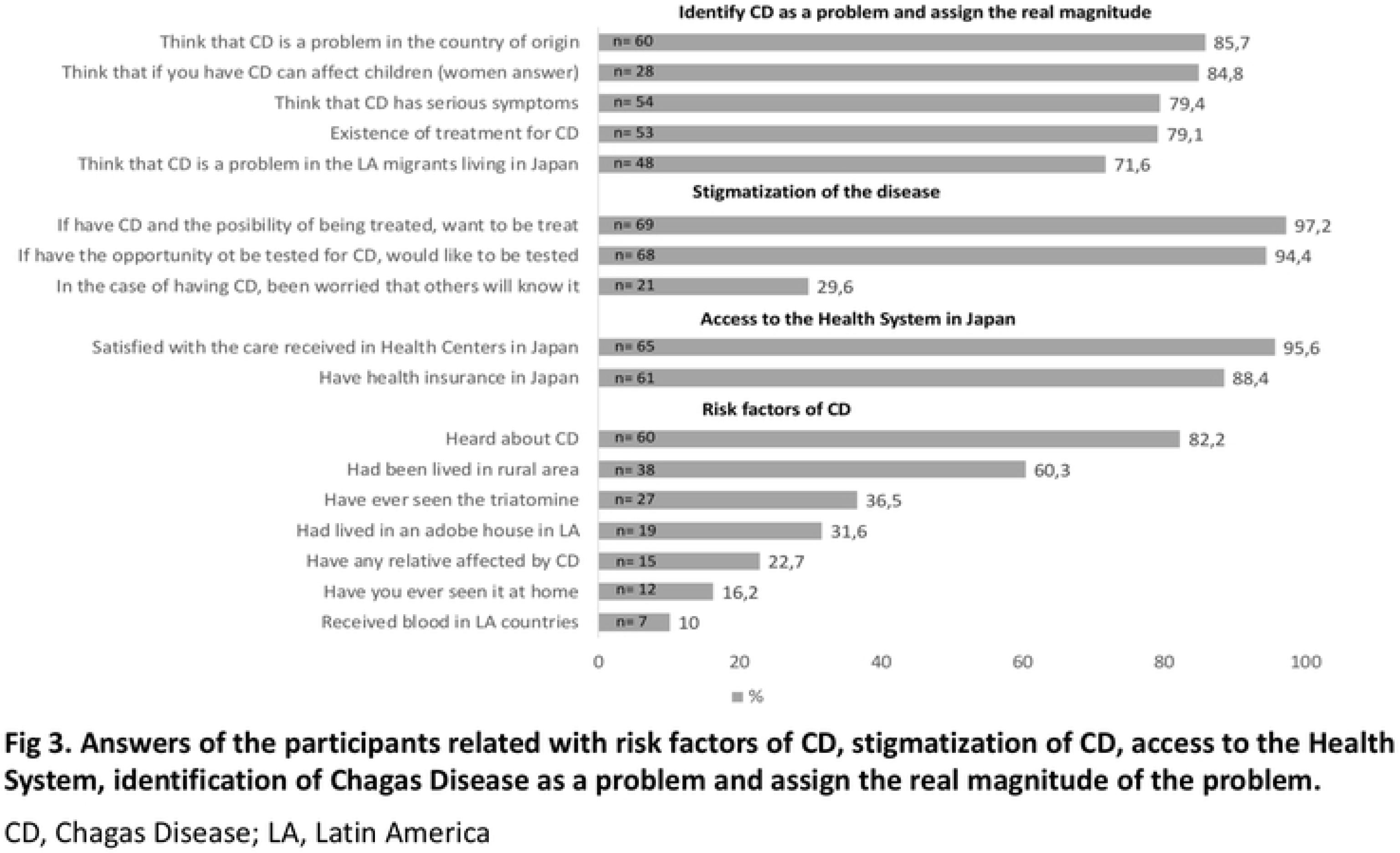
Answers of the participants related with risk factors of CD, stigmatization of CD, access to the Health System, identification of Chagas Disease as a problem and assign the real magnitude of the problem. CD, Chagas Disease; LA, Latin America

### Stigmatization of the disease

Most of the responders would like to be tested (94.4%) and treated (97.2%) for CD, if they had the opportunity. Nearly 30% would be worried if another person knew that they had CD (Fig 2).

### Access in terms of coverage and satisfaction to the health system

Most of the responders reported to have health insurance in Japan (88.6%) and in general, they were satisfied with the care received in health centers (95.6%) (Fig. 2).

### Knowledge of the participants

#### Baseline knowledge of the participants on Chagas disease

Our results showed that the total score of knowledge was low with the average at 6.7 ± 2.5. However, the participants could identify important points of the disease: epidemiology, transmission, symptomatology and treatment of CD. Most of the participants identified CD as a problem in LA countries (85.7%) and also in Japan (71.6%). Nearly 80% of them believed that CD was a severe disease (79.4%) (Fig. 2).

Most of the responders knew that CD was endemic in LA (n=27, 42.2%) with slightly difference in a second group that identified it worldwide (n=25, 39,0%). Nearly two thirds of the responders were familiar with the vector-borne transmission in first place (n=38). Most of the women (n=28, 84.8%) considered that the disease can affect their children if they are infected, just (n=12, 38.2%) of them answered vertical transmission as a possible route. Low knowledge about the oral transmission was reported between the participants. No misidentification of person to person route as hugs and kisses were answered by the responders.

A majority of the responders identify cardiac disease as a symptom of CD (n=53, 91.4%). Cardiac disease alone was the most popular answer with 58.6% of the answers (n=34). The next most answer was heart and digestive problems, both as part of the symptom of CD (n=17, 29.3%).

Two third of the responders considered a positive result to be interpreted as sick (n=40, 66,6%). Just 20% (n=12%) of the responders differentiated between infection and disease. Most of the responders (n=53, 79.1%) knew that there was treatment for CD and 69.2% (n=36) of them know that is not useful in severe cases.

#### Factors associated with better knowledge

Bivariable analysis showed that participants who had been living longer in Japan, heard about the disease, and had been tested for Chagas were significantly associated with the higher knowledge (P-value ≤ 0.2). Our multivariable logistic regression analysis indicated that longer living in Japan and prior test of Chagas were independently associated with the higher knowledge. Participants who lived in Japan for more than 10 years were 8 times (OR = 8.42, 95%CI: 1.56-48.62) more likely to be knowledgeable on CD than those who lived for shorter time. Participants who had been tested for CD were eleven times (OR. 11.32; 95%CI. 1.52-105.9) more likely to be knowledgeable on CD.

**Table 2.**
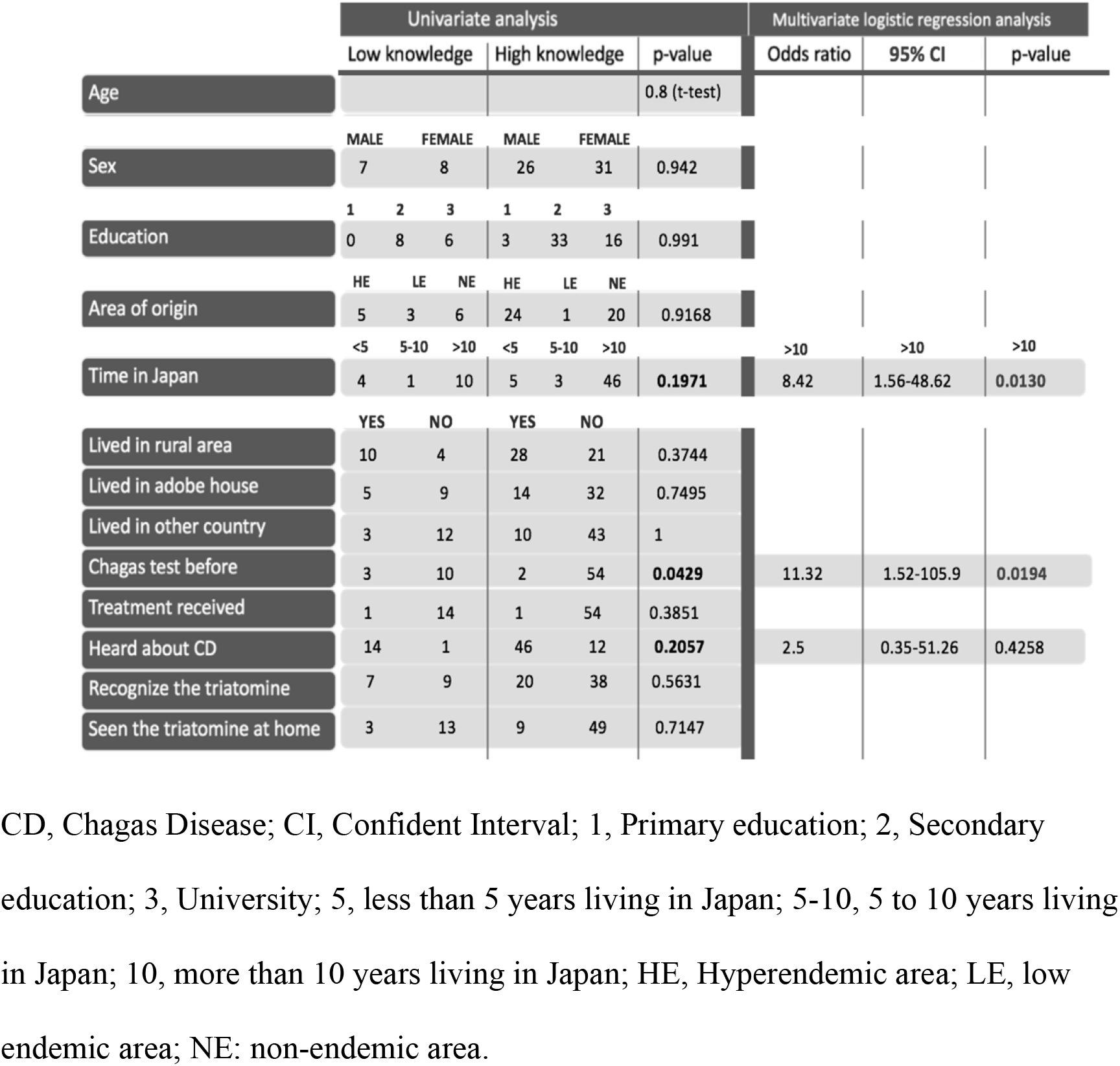
Univariate and multivariate analysis of factors associated with high knowledge of participants on the Chagas disease

#### Knowledge improvement after community activities

Sixty-one participants (out of 75) participated the CA (81.3%), while three people left the activity earlier and 11 arrived late to the venue. Among 61 participants participated in the CA, 50 (82%) of them performed the Questionnaire 1 and 2 (Fig. 3).

**Fig 3.**
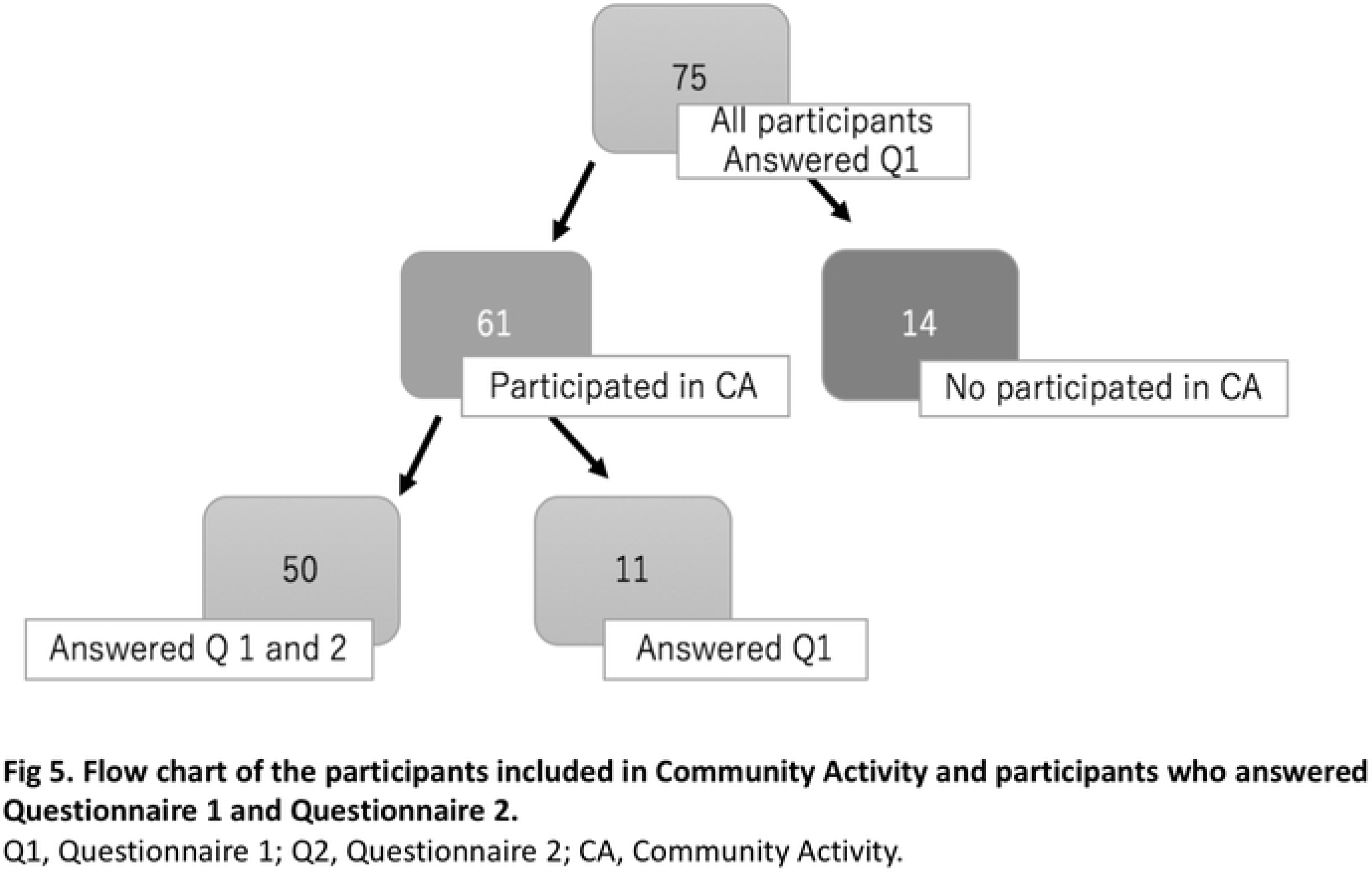
Flow chart of the participants included in Community Activity and participants who answered Questionnaire 1 and Questionnaire 2. Q1, Questionnaire 1; Q2, Questionnaire 2; CA, Community Activity.

Our paired sample t-test indicated an improvement in total knowledge of participants on CD (mean knowledge score: post-activities 9.8 ± 2.6 vs. pre-activities 6.7 ± 2.5, P-value < 0.0001; 95% CI: 2.32-3.84).

Before the community activities, only 32% of the participants (n=16) exhibited a high knowledge, however, 70% of the participants (n=35) had the high knowledge after the CA. As a result, more than 80% of the responders improved their punctuation after the activity (n=43, 86%). Just 2% (n=1) had worse punctuation after the CA and 12% (n=6) did not improved. Among the group that improved, the increase in points ranged from 0.5 to 10. Among the group that decreased their punctuation, the average reduction in point was 2.5.

### Qualitative analysis

#### Baseline knowledge

The responders were familiar with the disease and had prior knowledge about the epidemiology, vector-borne transmission and cardiac problems (S1 Table). Most of the participants were familiar with the word “Chagas”. They located the disease in their countries and their habits related to the high-attitude tropical areas. None was familiar with the flow of the disease into non-endemic areas (PC1 and 2).

The vector-borne transmission was most frequently identified as the only way of transmission. Some people who recognized blood transfusion as a way of transmission had the experience of a relative that was diagnosed during blood donation. Since they couldn’t donate blood, they associated the transmission with blood transfusion. Other routes as mother to child, organ transplantation or oral route were not considered before they were given information about it (PC3-4).

The participants identified the symptom of the disease mostly with cardiac problems. Digestive problem was less recognized but also was commented during FGD (PC5-7).

#### What is the first word that came to your mind when I say “Chagas”?

The most frequent answer was related to the feelings of fears, worries and memories. Fear. The fear was linked to CD for many participants. They expressed to have it for lack of information, for the severity of the disease and the belief fast death. Other participants manifested the fear of being ill out of their country of origin as a migrant (PC8-9).

Worries. Concerns were expressed for the transmission of the disease to their relatives. Women expressed more the concern when they learnt about the possibility of mother to child transmission (PC10-11).

Memories. The memories expressed during the FGD were related to the recent cases of CD that they experienced and were linked to feelings as fear or pain (PC12-13).

They also associated with the vector “vinchuca” and related with symptoms. The symptoms described were non-specific, related with pains or heart problems (PC14-15). Attitudes

### Attitudes

#### Normalization

Most of the participants never had been tested and they did not ask for the diagnosis test of Chagas. They represented a situation of normalization /naturalization of the disease (PC16). The fact of being sick or having some difficulties in the day life after the age of 50 was considered normal by the participants (PC17).

#### Stigmatization

During the FGD, the stigmatization of the disease did not become as a topic or commentary. When asked directly if they think that stigmas existed around CD or what they thought about the fact that some communities expressed stigma of CD, they denied the existence of stigmas. However, two participants thought differently. A participant expressed a situation of stigmatization, because of misbeliefs related to the route of transmission (PC18-19).

#### Resignation

Commentaries of resignation in front of a situation where death is inescapable had been described during FGD. This participant described the resignation in front a possibility to be infected linked to severe development of illness (PC20).

### Barriers

Even most of the participants were satisfied with the health system in Japan. Participants identify barriers related to the language, the cost of healthcare and familiarity with the health system in Japan that may affect the care seeking behavior of the participants. Also, their prioritization of the job as the main tool of support in a foreign country hindered them from getting care (PC21).

Language was described as one of the main barriers. They complained about the accuracy of the official translations and lack of intimacy (PC22).

They expressed prioritization other issues related with their migratory goals and not related to health if it was not necessary. Keeping the job position was one of the most important priorities (PC23-24).

The participants expressed problems in access to the health services. They were asked for more information if they wanted to make the diagnostic test of Chagas. They expressed the need to have the information in Spanish (P25).

### Participants proposals to overcome the actual problematic

Most of the participants claimed that more information should be provided. The participants considered that the spread of information in Spanish and by different media was an intervention that could improve the situation of CD in Japan. They proposed using the Internet as the most effective way to transmit information and offered a development of a website in Spanish with translated information. Furthermore, it is suggested to perform distribution of brochures or activities like the one held in our study in venues where LA migrant population tended to gather such as church, Latino meetings or health centers (PC26)

They expressed the importance to distribute information not just for people at risk, but also between the health sector personnel (PC27).

The participants highlighted the importance to have enough information before having the Chagas test. They remarked the need for accessible diagnosis, treatment and system of care (PC28-29).

## Discussion

To our knowledge, no study has analyzed the effect of educational activities in CD related to the knowledge of the participants. The fact that this activity included the participants as the main actors by sharing their knowledge and experiences, could have an important impact on the improvement of knowledge.

One of the factors that clearly had a significant influence on acquiring a high knowledge was being tested for CD in the past. People that had been tested before for CD had more information about the disease resulting a change in their care seeking behavior. One study conducted in Spain described that knowledge about the treatment efficacy could influence the decision to get the test of CD [16]. In our study, all participants who had been previously tested for CD were women, except one. This can be influenced by their reproductive role, as it was explained in other studies [17,11]. Living in Japan for more than 10 years also enhanced the knowledge probably due to the decrease of infested houses in the recent years in Bolivia and the urbanization of the population, along with the lack of prevention and care interventions [18,19]. This population migrated more than 10 years ago was more familiar with the vector. The urbanization of the population was proposed as a phenomenon that affected the awareness of the disease for its visualization in an area far from the traditional ruralism [20,11].

Similar to the result of other studies about CD knowledge of LA migrants, the baseline knowledge of our participants was low [21–24]. Due to high representation of Bolivian population recruited, results of our study were more similar to Blasco-Hernandez et al. study, conducted in Madrid in a group of Bolivian women [16]. The participants of our study had knowledge of CD including vector habitat, epidemiology, transmission by vector-borne and the cardiac problems associated with CD. The knowledge of blood transfusion transmission was achieved from previous experiences of having a relative diagnosed by CD during blood donation. Low knowledge of mother-to-child transmission was found in the women of our study. Therefore, this area required essential improvement to reduce new cases. Oral transmission was rarely considered as a way of transmission, which was indicated in another study [25]. Endemic countries have focused on reduction of the infestation by the vector; however, there are still lack of programs to control other ways of transmission, which can influence the awareness of the people.

Knowledge about symptoms and significance of a positive diagnosis were not achieved as baseline knowledge, as in the study of Blasco-Hernandez [16]. Many participants clarified the symptom as suffering for unspecific pain. The association with pain was also expressed in children in endemic area [25]. This may be influenced by the non-specificity of the CD clinic. The course of the disease is not well-known and it is irremediable associated with high mortality. The main difference with the study of Blasco-Hernandez et al., is that our participants were not CD-diagnosed population. Despite the prior knowledge about the disease, only 7.2% of the participants had been previously tested for CD. Most of the studies explained the low rate of diagnosis by the implication of socio-cultural factors on the representation of the disease. The most common factors are the stigmatization and the normalization of the disease [11, 12, 16, 17, 20, 21, 26, 27].

Dissemination of the information about CD was considered one of the most important strategies for improving the seeking for care. It was remarked by other studies conducted in non-endemic countries [24]. As in other studies, religious meetings, social association and graphic materials was proposed for disseminate information. However, it was proposed that the Internet was the most effective way to provide the information. Most of the population at risk of CD are highly stigmatized and have experienced discrimination. However, our participants showed a low level of stigmatization. One of the possible explanations is that most of them migrated at the age of 30 years-old. They remembered cases of CD of old relatives, but not proximal generation. This might make them less familiar to the reality of CD in their socio-cultural context and feel less vulnerable. Another possible explanation is that most of our participants were not diagnosed with CD, in contrast with other studies that shows a high level of stigmatization [28]. As in most of the populations analyzed for CD, our participants showed a high level of normalization/naturalization of the disease. The un-specificity of symptoms and the lack of impact on day-to-day activities as described in many studies contributes to the normalization of the disease. Also, similar to the study by Blasco-Hernandez et al, the participants of our study were normalized to be ill at the age of 50. The low social-relevance of CD was influenced by the low expected age of life in Bolivia in the last decade. However, this data is changing nowadays [28].

The representation of the disease for the participants were similar to the one described previous studies. They presented a disease caused by the “vinchuca”, an insect that affects the heart and leads in a high mortality. The reactions and associations with CD after this representation, were predominated by feelings of fears, worries, and memories of affected relatives. As previously described, this emotional burden led to the attitudes of resignation until the irremediable in some participants. The long period asymptomatic and a late diagnosis contributed to the maintenance of this representation. This study was conducted in Japan, a country that has an important gap in epidemiological data for CD. The literature of CD in Japan is limited to cases report and one study of prevalence in the Brazilian population [29]. This is the first study to analyze the LA migrant population living in Japan, as a population at risk of CD. However, most of participants were Bolivians. This representation could cause a recruitment bias, since the Embassy of Bolivia was the only embassy to be part of the study. Santa Cruz was the most popular area of origin. This was expected because the migration in the Japanese citizens to Bolivia after the second world war II. They were installed in the district of Santa Cruz, a high endemic area. Two Japanese colonies near the city of Santa Cruz were founded in the 50s, “Okinawa Colony” and “San Juan of Yapacaní Colony”.

As migrant population, several barriers were identified in the process of seeking for health care in Japan. Preconceptions conceived in the country of origin may act as a barrier in the host country. The barriers found in Japan included language barrier, migratory process, and difficulties to access the health care system. These results are similar to the barriers experienced by the LA migrant population in other non-endemic countries [30, 11, 17, 31].

Barriers of accessing to the health system for migrants are highly documented in different studies. The lack of adaptability to facilitate the access of migrant population is constant in host countries [17, 21, 31, 32, 33]. In our study, most of the participants had not known where to seek for care in case of desire to be tested for CD. This can be responsible by the organization of the Health System in Japan, where the role of family doctor is uncommon, which adds difficulties for the people at risk to find the integrative care for a disease such as Chagas.

On the other hand, there has still not been study conducted in Japan to analyze the knowledge of Japanese doctors about tropical diseases. However, this knowledge is expected to be low as happening in most of the non-endemic countries [34].

The understanding of the implications of social and cultural factors described in care seeking behavior of the migrant population is a key for designing policies, control and preventive interventions. These interventions should avoid the traditional specific disease-centered due to the low impact on the population [11].

One of the main strengths of this study is the transnational approach. Most of the participants were not tested for CD before. It provides a naïve vision of the knowledge, conceptions and representations of the disease in the population at risk. The study was conducted in Spanish by a native speaker to ensure the cultural competence and the feasibility of the data in the qualitative part.

We did not identify the most successful gathering method. However, most of the participants coming to the venue had some other issues to resolve with the Embassy of Bolivia. It leads in a low representation of other LA countries with high representation in Japan such as Brazil, Peru and Argentina. This can lead to a possible recruitment bias. Also, the activities were held on Saturday, which is a working day in Japan. The questionnaires used in this study were mostly based on previous studies. However, it has not been validated. This activity demonstrates an increase of knowledge. However, it could have a memory bias and we did not identify the impact of the impact of knowledge on care seeking behavior. The population that did not participate in the community activity were much younger than the participated ones. Improvement should be designed to reach to young population. A virtual format in networks, a more actual way of communication or approaches in venues which are more familiar to teenagers such as schools or activity clubs should be considered.

In conclusion, educational activities with integral approach are useful to increase the knowledge of CD. This activity brings the possibility to explore not only the knowledge but also the characteristic, experiences, opinions and needs of those at risk. This information is essential in order to guide the efforts to improve the CD problem, considering the people at risk as part of the improvement and development. However, the effectiveness of this activity should be evaluated in different geographical areas. Longitudinal research will bring more information on how the knowledge acquired by integral activities influences the seeking behaviour in Chagas disease.

## Supporting information

**S1 Table. Qualitative analysis commentaries** Each participants commentaries (PC) was numbered as PC1-28 shown in the supplement table (S1).

## Acknowledgements

Authors wish thank to all the members of Bolivian Embassy for the important and essential collaboration in this project. I would like to appreciate the generous support provided by the leaders of the LA community in the different cities: Mr. Miguel Saucedo in Oizumi, Mrs. Vania Sikujara in Suzuka, the pastor Mr. Jaime Teruya in Hadano and Mr. Jorge Añez in Nagoya. Roxana Oshiro for the great collaboration in the expansion of the activity by the media. Finally, we would like to thank to the participants for sharing their experience to the community.

